# Longitudinal metatranscriptomic sequencing of Southern California wastewater representing 16 million people from August 2020-21 reveals widespread transcription of antibiotic resistance genes

**DOI:** 10.1101/2022.08.02.502560

**Authors:** Jason A. Rothman, Andrew Saghir, Seung-Ah Chung, Nicholas Boyajian, Thao Dinh, Jinwoo Kim, Jordan Oval, Vivek Sharavanan, Courtney York, Amity G. Zimmer-Faust, Kylie Langlois, Joshua A. Steele, John F. Griffith, Katrine L. Whiteson

## Abstract

Municipal wastewater provides a representative sample of human fecal waste across a catchment area and contains a wide diversity of microbes. Sequencing wastewater samples provides information about human-associated and medically-important microbial populations, and may be useful to assay disease prevalence and antimicrobial resistance (AMR).

Here, we present a study in which we used untargeted metatranscriptomic sequencing on RNA extracted from 275 sewage influent samples obtained from eight wastewater treatment plants (WTPs) representing approximately 16 million people in Southern California between August 2020 – August 2021. We characterized bacterial and viral transcripts, assessed metabolic pathway activity, and identified over 2,000 AMR genes/variants across all samples. Because we did not deplete ribosomal RNA, we have a unique window into AMR carried as ribosomal mutants. We show that AMR diversity varied between WTPs and that the relative abundance of many individual AMR genes/variants increased over time and may be connected to antibiotic use during the COVID-19 pandemic. Similarly, we detected transcripts mapping to human pathogenic bacteria and viruses suggesting RNA sequencing is a powerful tool for wastewater-based epidemiology and that there are geographical signatures to microbial transcription. We captured the transcription of gene pathways common to bacterial cell processes, including central carbon metabolism, nucleotide synthesis/salvage, and amino acid biosynthesis. We also posit that due to the ubiquity of many viruses and bacteria in wastewater, new biological targets for microbial water quality assessment can be developed.

To the best of our knowledge, our study provides the most complete longitudinal metatranscriptomic analysis of a large population’s wastewater to date and demonstrates our ability to monitor the presence and activity of microbes in complex samples. By sequencing RNA, we can track the relative abundance of expressed AMR genes/variants and metabolic pathways, increasing our understanding of AMR activity across large human populations and sewer sheds.

## 1. Introduction

Wastewater harbors a wide diversity of microorganisms and represents the collective waste of human activity across a sewershed (Newton and McClary, 2019). Over 300 km^3^ of wastewater is produced globally, of which most is channeled into wastewater treatment plants (WTPs) for biological and chemical processing (Lu et al., 2018). As a heterogenous mixture, wastewater has been shown to contain microbial communities that vary depending on sampling location, time of year, industry, agriculture, and the health of the served human population (Cantalupo et al., 2011; Edwards et al., 2019; McLellan et al., 2010; Symonds et al., 2009; Wu et al., 2019). As a result, the microbial water quality of wastewater can be a useful indicator of an area’s biological contamination, with outbreaks of several diseases corresponding to increased wastewater titers of pathogenic etiological agents (Hellmér et al., 2014; Manor et al., 1999; Rothman et al., 2021; Wu et al., 2020). The microbial ecology of wastewater is an important topic, with many studies characterizing the microbes present through culturing, PCR- and sequencing-based methods, and generally rely on targeting specific pathogens or metagenomic shotgun DNA sequencing (Hubeny et al., 2022; Jankowski et al., 2022; Kitajima et al., 2018; Martínez-Puchol et al., 2020). While useful, these studies are unable to capture microbial transcription, which provides information about active microbial processes, instead of the genomic potential of wastewater. Moreover, as many important human and crop/livestock pathogens are RNA viruses (Amoah et al., 2020; Bibby and Peccia, 2013; Symonds et al., 2009), we can monitor the presence and spread of *Ribovira* through untargeted metatranscriptomics. Wastewater RNA sequencing can uncover active microbial interactions and metabolic networks, which may inform us of the public and environmental health of the areas served by a given sewage system (Brumfield et al., 2022; Crits-Christoph et al., 2021; Li et al., 2022; Rothman et al., 2021).

Wastewater-based epidemiology (WBE) can inform public health about the presence of pathogens in a population without needing to test individuals in healthcare settings (Bivins et al., 2020; Sims and Kasprzyk-Hordern, 2020). Health agencies have used and WBE to detect the presence of human pathogens such as norovirus, polio, SARS coronaviruses, and a variety of bacteria and protists (Hellmér et al., 2014; Manor et al., 1999; Rothman et al., 2021; Wu et al., 2020). For example, WBE has been heavily used to track and monitor the abundance and spread of SARS-CoV-2 during the ongoing COVID-19 pandemic at various population levels (Achak et al., 2021; Karthikeyan et al., 2021; Nemudryi et al., 2020; Peccia et al., 2020; Rothman et al., 2021; Wu et al., 2020). Furthermore, as disease case counts change longitudinally, multiple time points and RNA sequencing are useful to track not only the presence, but the activity of microorganisms which may provide additional information about pathogens over longer time periods (Faust et al., 2015; Joseph et al., 2019; Marcelino et al., 2019; Nemudryi et al., 2020). Lastly, by broadly sequencing RNA, we may be able to discover new targets for microbial water quality assays in order to detect and monitor for sewage contamination of the environment and water sources (Cao et al., 2015; Farkas et al., 2019; Jiang et al., 2022; Kitajima et al., 2018; Zimmer-Faust et al., 2021).

Antimicrobial resistance (AMR) is a worldwide concern that inhibits effective treatment of disease and increases healthcare burden and morbidity of infections (World Health Organization, 2021). Wastewater contains a complex diversity of AMR genes, which allows for horizontal gene transfer (HGT) between antimicrobial resistant organisms and those species or strains that are currently susceptible to antimicrobial therapies (Joseph et al., 2019; Ju et al., 2019; Sims and Kasprzyk-Hordern, 2020). As AMR and HGT are important to monitor for public and agricultural health, many studies have employed sequencing and targeted PCR-based technologies to assay the AMR genomic content of wastewater (Karkman et al., 2018). While useful, these studies typically rely on DNA-based technologies which cannot measure the transcriptional activity of these genes or the organisms that harbor AMR, and may better indicate the severity and abundance of antimicrobial resistant infections across a population (de Nies et al., 2021; Ju et al., 2019; Marcelino et al., 2019). By employing RNA-sequencing, we are better able to understand the disease ecology and AMR activity of wastewater-inhabiting organisms and those deposited through the waste stream, and the specific mutations that cause AMR (Alcock et al., 2020). Lastly, through careful sampling, changes in AMR transcription can be tracked over time, likely providing finer-scale information about the severity and seasonality of AMR infections during the ongoing COVID-19 pandemic (Langford et al., 2020; Rose et al., 2021).

Studying wastewater microbial ecology and tracking the activity of disease-associated microbes and AMR is vital to public health and environmental monitoring. In this study, we used metatranscriptomic sequencing to characterize the RNA world of 275 samples across eight wastewater treatment plants (WTPs) representing approximately 16 million people across Southern California. We investigated several lines of inquiry: First, what is the transcriptomic diversity of microorganisms in Southern California wastewater, and does it vary longitudinally? Second, what AMR genes are being actively transcribed in wastewater? Third, are there conserved biochemical pathways across wastewater, and does this metabolic potential vary? Lastly, are there largescale patterns of microbial transcription in Southern California’s wastewater, and is there a temporal component to any of these patterns?

## 2. Materials and Methods

### 2.1 Sample collection

We previously reported the sample collection and handling procedure in Rothman et al 2021 (Rothman et al., 2021), and note that the viromes of 94 samples were previously reported in that study. Briefly, we collected 275 1-liter 24-hour composite influent wastewater samples by autosampler at eight WTPs across Southern California between August 2020 – August 2021 (Table 1). We aliquoted and stored 50 mL of sample at 4 °C until RNA extraction.

**Table 1.**
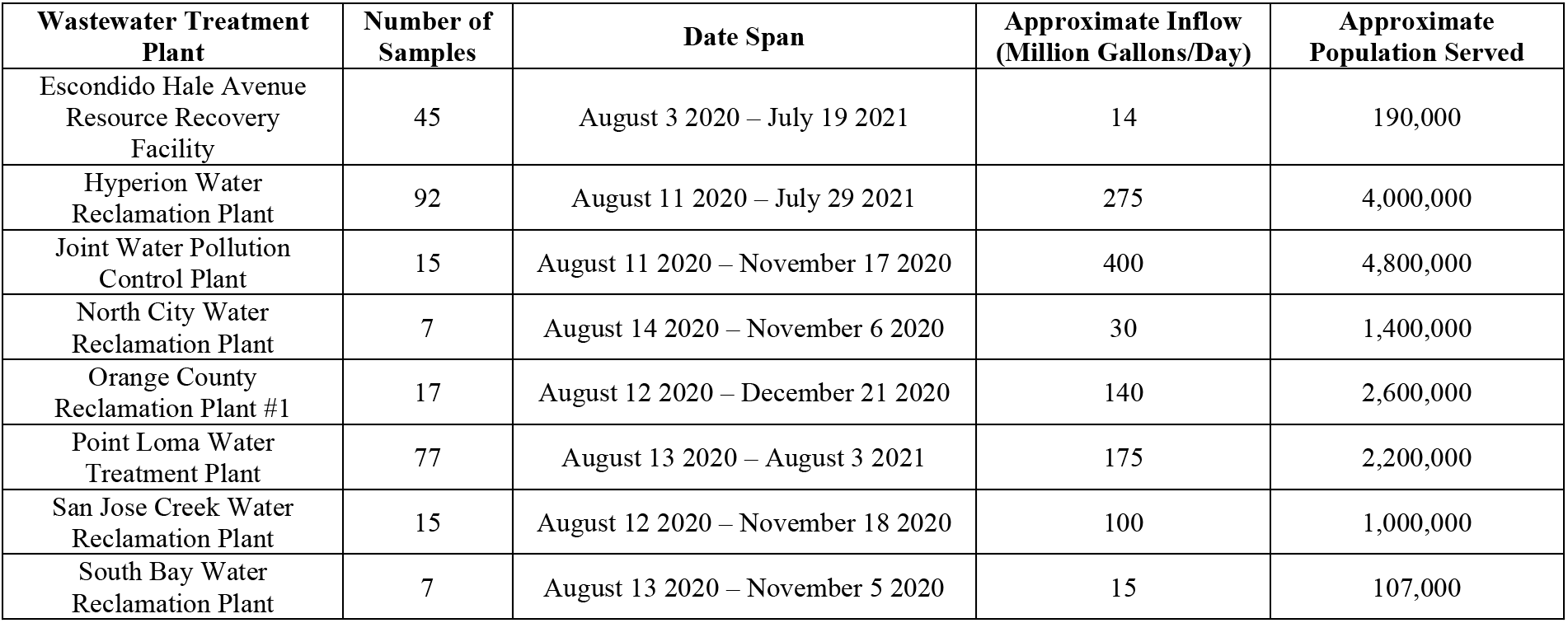
Descriptions of the experiment sampling scheme and relevant information about each WTP.

| **Wastewater Treatment Plant** | **Number of Samples** | **Date Span** | **Approximate Inflow (Million Gallons/Day)** | **Approximate Population Served** |
| --- | --- | --- | --- | --- |
| Escondido Hale Avenue Resource Recovery Facility | 45 | August 3 2020 – July 19 2021 | 14 | 190,000 |
| Hyperion Water Reclamation Plant | 92 | August 11 2020 – July 29 2021 | 275 | 4,000,000 |
| Joint Water Pollution Control Plant | 15 | August 11 2020 – November 17 2020 | 400 | 4,800,000 |
| North City Water Reclamation Plant | 7 | August 14 2020 – November 6 2020 | 30 | 1,400,000 |
| Orange County Reclamation Plant #1 | 17 | August 12 2020 – December 21 2020 | 140 | 2,600,000 |
| Point Loma Water Treatment Plant | 77 | August 13 2020 – August 3 2021 | 175 | 2,200,000 |
| San Jose Creek Water Reclamation Plant | 15 | August 12 2020 – November 18 2020 | 100 | 1,000,000 |
| South Bay Water Reclamation Plant | 7 | August 13 2020 – November 5 2020 | 15 | 107,000 |

### 2.2 RNA extraction and sequencing library preparation

We used a protocol based on Crits-Christoph 2021 (Crits-Christoph et al., 2021) and Wu et al 2020 (Wu et al., 2020), in which we pasteurized 50 mL of influent wastewater in a 65 °C water bath for 90 minutes, then filtered samples through a sterile 0.22-μM filter (VWR, Radnor, PA). We then centrifuged the sample at 3,000 xg through 10-kDa Amicon filters (MilliporeSigma, Burlington, MA) and stored the concentrate at −80 °C until RNA extraction. We then used an Invitrogen PureLink RNA Mini Kit with added DNase step (Invitrogen, Waltham, MA) following the manufacturer’s instructions to extract RNA and stored the resulting RNA at −80 °C until library preparation.

The University of California Irvine Genomics High Throughput Facility (GHTF) handled all library preparation steps. Briefly, the GHTF used the Illumina RNA Prep for Enrichment kit (Illumina, San Diego, CA) on each RNA sample, then sequenced the paired end libraries as 2 x 100bp or 2 x 150 bp (supplemental file SF1) with an S4 300 cycle kit on an Illumina NovaSeq 6000 over four batches.

### 2.3 Bioinformatics and data processing

We received the data from the GHTF as demultiplexed FASTQ files and used the UCI High Performance Community Computing Cluster for data processing. We used BBTools “bbduk” (Bushnell, 2014) to remove adapters, low-quality bases, and primers, then removed PCR duplicates with BBTools “dedupe.” After deduplication, we removed reads mapping to the human genome (hg38) with Bowtie2 (Langmead and Salzberg, 2012), then used Kraken2 (Wood et al., 2019) and Bracken (Lu et al., 2017) databases built with the NCBI RefSeq database of bacteria, archaea, and viruses (January 2021), to taxonomically classify reads. We then tabulated these reads and used these tables for downstream diversity analysis (supplemental file 2).

For community analyses, we normalized the transcript reads into within-sample relative abundances in R, removed reads corresponding to less than 0.01% relative abundance, then calculated Shannon Diversity indices and Bray-Curtis dissimilarity matrices with the R package “vegan” (Oksanen et al., 2017). We generated nonmetric multidimensional scaling (NMDS) ordinations, then tested the diversity metrics for significant differences with Kruskal-Wallis tests (alpha diversity) and Adonis PERMANOVA (beta diversity) with “vegan.” We assessed the relationship of diversity with time with linear mixed effects models (lmer) in the R package “lmerTest” using WTP and sequencing batch as random effects (Kuznetsova et al., 2017), and plotted all diversity analyses with “ggplot2” (Wickham, 2009), “ggrepel” (Slowikowski, 2018), and “patchwork” (Pedersen, 2020). Because we collected samples from Escondido, Hyperion, and Point Loma for a much longer period of time than the other WTPs, we ran the above analyses two ways: all WTPs together from August – November 2020, and Escondido, Hyperion, and Point Loma samples for the full year separately.

We used HUMAnN3 (Beghini et al., 2021) with default settings to assign functional gene pathway annotations to reads using the UniRef90 (Suzek et al., 2015) and Metacyc (Caspi et al., 2020) databases. We also used RGI (the Resistance Gene Identifier) and the CARD and WildCARD databases (Alcock et al., 2020) to assign predicted antimicrobial resistance ontology identities (AROs) to the reads, then normalized all pathway abundances and AMR gene identities to transcripts per million (TPM). We compared microbial abundances, pathway abundances, and AMR gene abundances at greater than 0.01% relative abundance and present in 50% of samples between WTPs with ANCOM2.1 using sample collection month as an adjustment for covariates and sequencing batch as a random effect in the ANCOM models. We then plotted log_10_ transformed counts of significantly differentially abundant viruses, bacterial genera, and AMR genes on a heatmap allowing the taxa to cluster with the Ward D2 algorithm with the R package “pheatmap” (Kolde, 2019). We used MaAsLin2 (Mallick et al., 2021) for longitudinal analyses of the above-mentioned variables, and included WTP and sequencing batch as random effects in the models, and we adjusted ANCOM and MaAsLin2 statistical tests for multiple comparisons with the Benjamini-Hochberg correction. We report the linear model coefficient with time of MaAsLin2 analyses on each plot and refer to Zenodo (doi.org/10.5281/zenodo.6829029) (Rothman et al., 2022) for individual scatterplots.

### 2.4 Data and code availability

Representative analyses scripts and code are available at github.com/jasonarothman/wastewater_metatranscriptomics_socal_aug20_aug21 and raw sequencing files have been deposited at the NCBI Sequence Read Archive under accession numbers PRJNA729801. Data tables containing taxa abundances, HUMAnN3 pathway annotations, and RGI assigned predicted antimicrobial resistance ontology identities are available as a Dryad dataset (https://doi.org/10.7280/D11Q30) (Rothman et al., 2022)

## 3. Results

### 3.1 Library statistics and microbial sample composition

We obtained a total of 4,336,566,730 quality-filtered, nonhuman, paired-end reads across 275 samples from eight WTPs (average: 15,769,334 reads per sample, range: 1,039,430 – 88,651,858). With Kraken2, we classified an average of 55.0% of our reads (range 8.7 – 83.5%), of which an average of 48.1% (range 7.0 – 83.0%) were bacterial, 0.2% were archaeal (range 0.02 – 3.8%), and 5.9% (range 0.03 – 38.3%) were viral (Fig. 2). Due to the low relative abundance of archaea and known questionable classification accuracy, we chose to focus on bacteria and viruses for diversity analyses.

**Figure 1.**
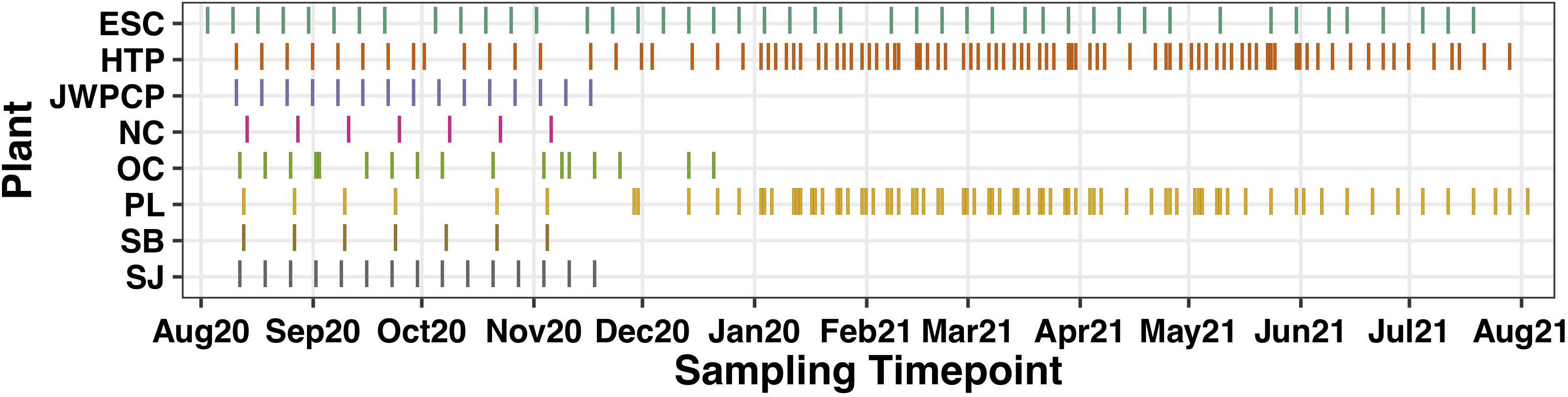
Diagram indicating the date ranges of samples separated by wastewater treatment plan and month. Y-axis codes correspond to abbreviated WTP names: ESC = Escondido Hale Avenue Resource Recovery Facility, HTP = Hyperion Water Reclamation Plant, JWPCP = Joint Water Pollution Control Plant, NC = North City Water Reclamation Plant, OC = Orange County Reclamation Plant #1, PL = Point Loma Water Treatment Plant, SJ = San Jose Creek Water Reclamation Plant, and SB = South Bay Water Reclamation Plant.

**Figure 2.**
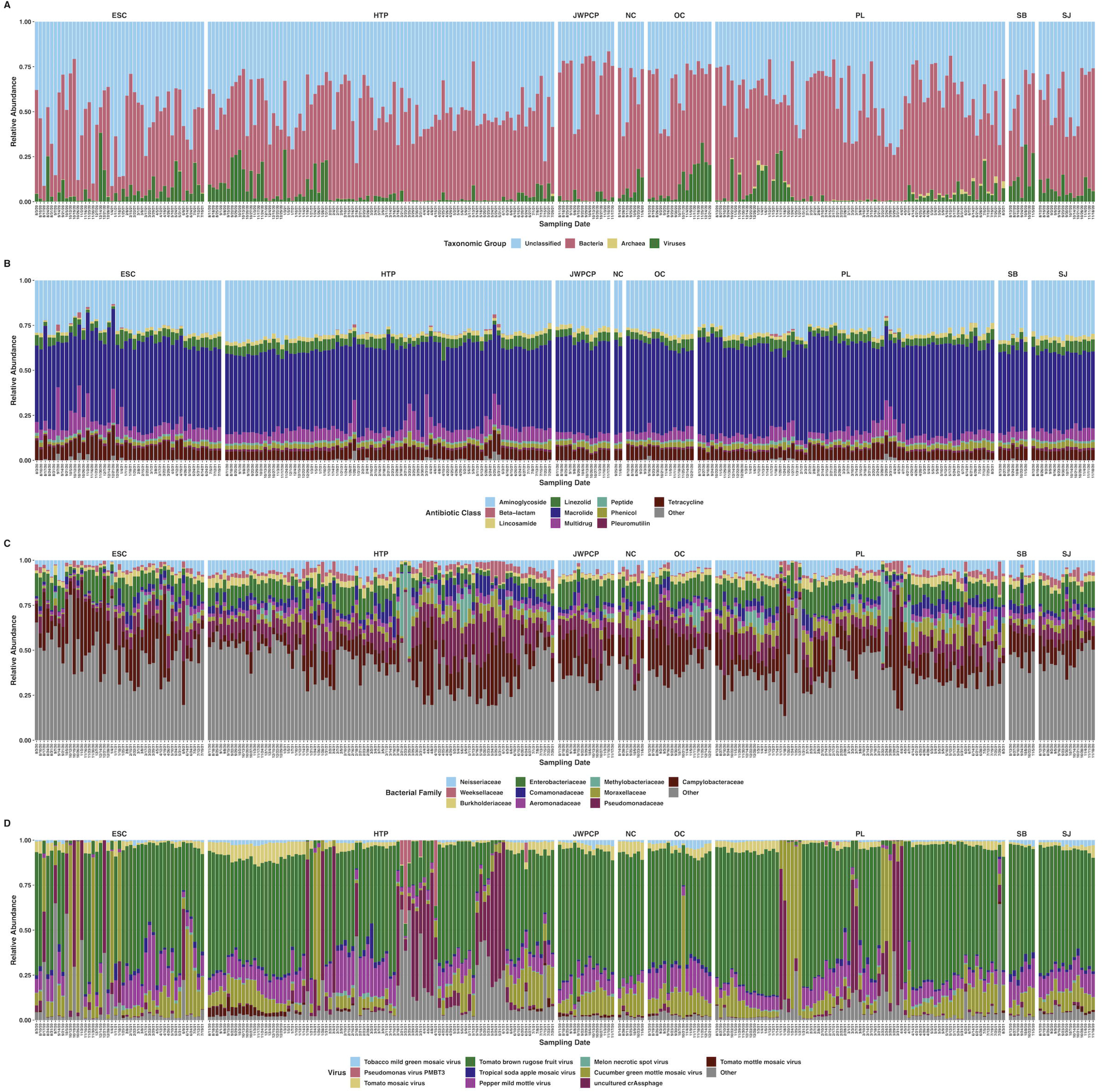
Stacked bar plots showing the relative abundances of RNA reads mapping to A) unclassified taxonomic ranks, bacteria, viruses, and archaea; B) AMR genes separated by the ten most abundant antibiotic classes each gene confers resistance to plus all others; C) ten most abundant bacterial families plus all others; D) ten most abundant viral species plus all others. All plots are faceted by WTP and labeled with sampling date.

We detected transcripts from a total of 6,449 bacterial and 6,888 viral species across all samples, however due to the likelihood of the taxonomic classifier reporting spurious species, we removed species accounting for < 0.01% average relative abundance within each domain. This filtering left us with 935 bacterial and 134 viral species present, which we used for downstream analyses. We also tabulated 245 bacterial families present in the same fashion as above. Because we had an uneven longitudinal distribution of samples, we analyzed diversity, differential abundance, and longitudinal relationships in two ways: First, samples where we had all eight WTPs were analyzed together representing N = 98, covering the months of August – November 2020. Second, we analyzed samples from Escondido, Hyperion, and Point Loma WTPs, where we had an entire year of sampling (N = 214), covering August 2020 – August 2021.

### 3.2 Antimicrobial resistance transcription

We detected transcripts matching 2,128 unique antibiotic resistance ontology identifiers (AROs) through use of RGI and the CARD database (Fig. 2, Dryad: https://doi.org/10.7280/D11Q30). AMR alpha diversity between August – November 2020 significantly differed between WTPs (H_(7)_ = 33.7, P < 0.001), but not over time (t = −0.3, P = 0.74). AMR beta diversity during this time only differed between WTPs (P < 0.001, R^2^ = 0.43), and not by month (P = 0.08, R^2^ = 0.03), an interaction of WTP and month (P = 0.10, R^2^ = 0.16), or sequencing batch (P = 0.05, R^2^ = 0.02), and slightly changed over time (t = −2.3, P = 0.03) (Fig. 3). Several AMR transcripts were differentially abundant between WTPs, and for easier discrimination between the categories, we separated them into ribosomal RNA mutations and non-rRNA AMR genes: 29 rRNA AMR mutants (W > 88, P_adj_ < 0.05) and 17 non-rRNA genes (W > 140, P_adj_ < 0.05) differed between WTPs (Fig. 3, supplemental file SF2).

**Figure 3.**
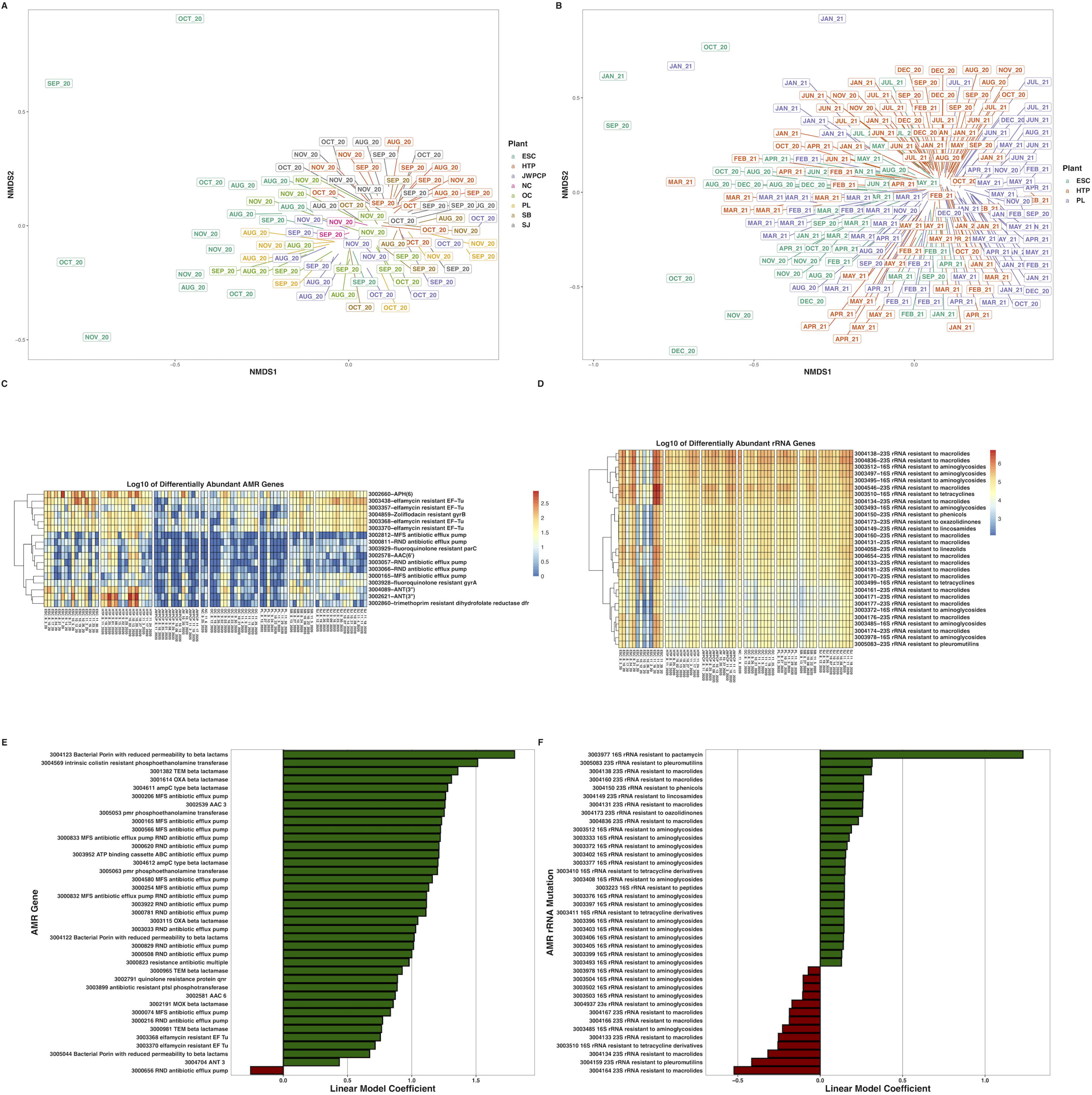
Nonmetric multidimensional scaling ordination of Bray-Curtis dissimilarities of AMR genes at greater than 0.01% relative abundance across A) all WTPs August – November 2020, and B) ESC, HTP, and PL across August 2020 – August 2021. C) Heatmaps of the log_10_-transformed counts of differentially abundant non-rRNA AMR genes across all WTPs August – November 2020, and D) rRNA gene mutations conferring resistance to antimicrobials. Hierarchal clustering of genes in each heatmap is through the Ward D2 algorithm. E) Bar plots indicating the non-RNA AMR genes across ESC, HTP, and PL that changed over time and F) AMR rRNA gene mutations. X-axes denote the linear model coefficient of each gene’s relationship to time.

When considering the entire year, AMR alpha diversity differed between WTPs (H_(2)_ = 28.6, P < 0.001), but not over time (t = −0.6, P = 0.68). AMR beta diversity differed between WTPs (P < 0.001, R^2^ = 0.13), month (P < 0.001, R^2^ = 0.16), an interaction between WTP and month (P = 0.007, R^2^ = 0.12), with significant batch effects (P < 0.001, R^2^ = 0.06), and again, changed over time (t = 3.3, P = 0.001) (Fig. 3). We considered AMR transcripts from rRNA genes and non-ribosomal genes separately as above. For rRNA genes, we found that 26 positively and 13 negatively correlated with time (P_adj_ < 0.05), while 45 did not, and for non-ribosomal genes, 38 positively and 1 negatively changed over time (P_adj_ < 0.05), while 256 did not change significantly (Fig. 3, supplemental file SF3, Zenodo: doi.org/10.5281/zenodo.6829029).

### 3.3 Bacterial transcriptional ecology

We found that the top ten most proportionally abundant bacterial families represented an average of 58.6% (range 17.7 – 82.3%) of bacterial transcripts. These families (in descending average proportional abundance) were: Campylobacteraceae, Pseudomonadaceae, Enterobacteriaceae, Neisseriaceae, Moraxellaceae, Comamonadaceae, Burkholderiaceae, Aeromonadaceae, Weeksellaceae, and Methylobacteriaceae (Fig. 2).

From August – November 2020, bacterial transcript alpha diversity significantly differed between WTP (H_(7)_ = 55.5, P < 0.001), but not over time (t = −1.3, P = 0.22). Bacterial beta diversity was significantly different across WTPs (P < 0.001, R^2^ = 0.30) and month (P < 0.001, R^2^ = 0.09), with no interaction between WTP and month (P = 0.13, R^2^ = 0.16), was affected by sequencing batch (P < 0.001, R^2^ = 0.03), and changed over time (t = −2.4, P = 0.02). We also found that transcripts from 222/564 bacterial genera were significantly differentially abundant between WTPs during this time period (W > 507, P_adj_ < 0.05, Fig. 4, supplemental file SF2).

**Figure 4.**
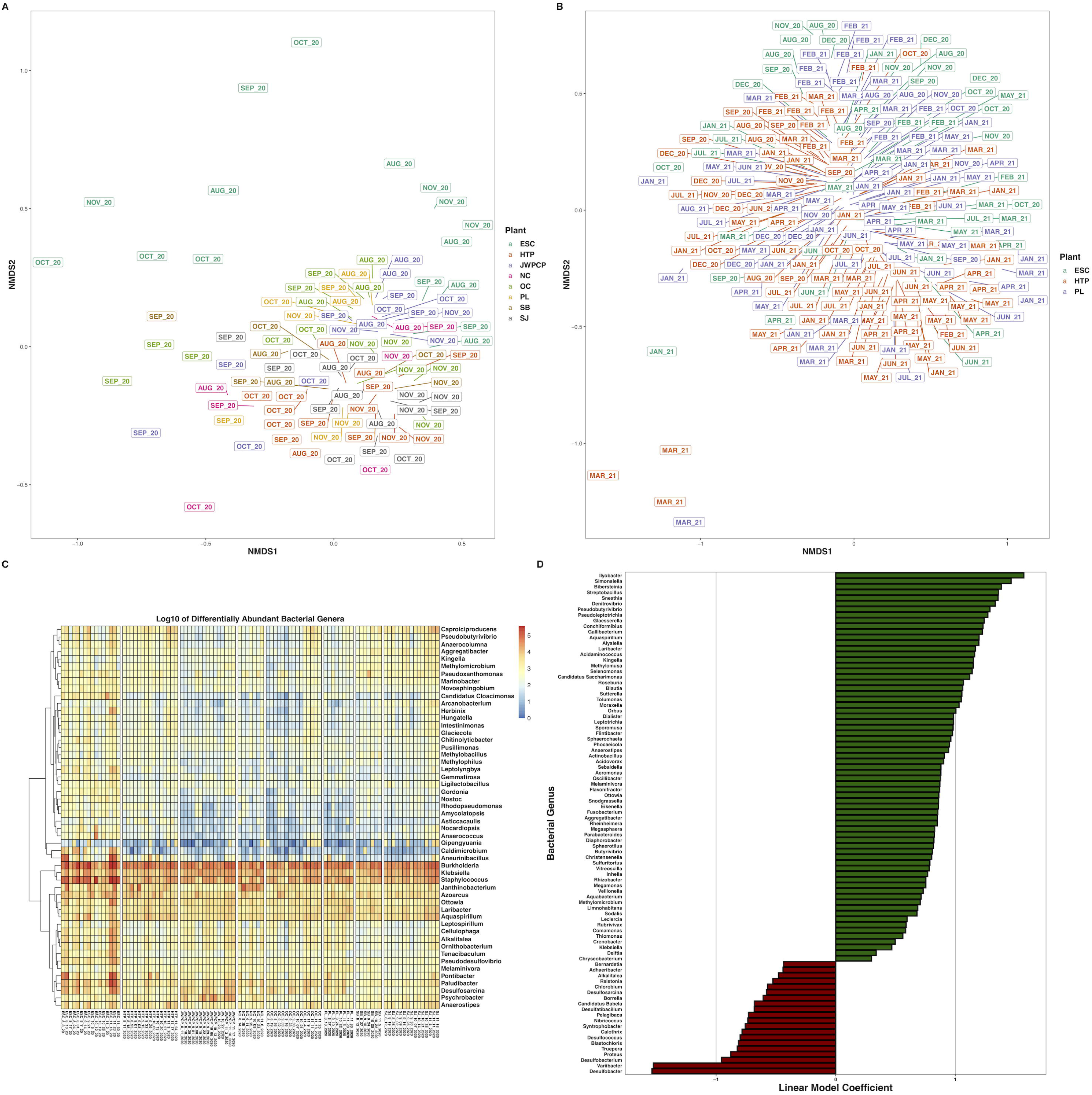
Nonmetric multidimensional scaling ordination of Bray-Curtis dissimilarities of bacterial species at greater than 0.01% relative abundance across A) all WTPs August – November 2020, and B) ESC, HTP, and PL across August 2020 – August 2021. C) Heatmap of the log_10_-transformed counts of differentially abundant bacterial genera at greater than 0.1% relative abundance across all WTPs August – November 2020. D) Bar plots indicating the bacterial genera across ESC, HTP, and PL that changed over time (only genera with a P_adj_ < 0.001 shown). X-axes denote the linear model coefficient of each genus’s relationship to time.

Across the entire year, alpha diversity was not different between WTPs (H_(2)_ = 1.1, P = 0.59), and did not differ over time (t = 1.6, P = 0.12). Beta diversity differed between WTP (P < 0.001, R^2^ = 0.07), month (P < 0.001, R^2^ = 0.20), and the interaction of WTP and month (P = 0.002, R^2^ = 0.11) with significant batch effects (P < 0.001, R^2^ = 0.06), and over time as a continuous variable (t = 4.8, P < 0.001). We tracked the transcription of bacterial genera across the year, and found that 172 genera increased, 63 genera decreased, and 295 did not change significantly over time (Fig. 4, supplemental file SF3, Zenodo: doi.org/10.5281/zenodo.6829029).

### 3.4 Viral ecology

We did not group viruses by family because of the dominance of Virgaviridae, and instead report summary statistics of the ten most proportionally abundant viral species as this provides more information. These viruses represented an average proportional viral abundance of 92.4% (range 33.1 - 99.5%; in descending average proportional abundance): Tomato brown rugose fruit virus, Cucumber green mottle mosaic virus, Pepper mild mottle virus, crAssphage, Tomato mosaic virus, Tropical soda apple mosaic virus, Tobacco mild green mosaic virus, Tomato mottle mosaic virus, Melon necrotic spot virus, and Pseudomonas virus PMBT3 (Fig. 2).

Over August – November 2020, viral alpha diversity differed between WTPs (H_(7)_ = 35.1, P < 0.001), but not over time (t = −0.57, P = 0.58). Beta diversity differed between WTPs (P < 0.001, R^2^ = 0.31), by month (P = 0.003, R^2^ = 0.07), but not by an interaction between WTP and month (P = 0.69, R^2^ = 0.13), by batch (P = 0.07, R^2^ = 0.02), or over time (t = −1.6, P = 0.11). During this time period, only 11 viruses were differentially abundant between WTPs (Fig. 5, supplemental file SF2).

**Figure 5.**
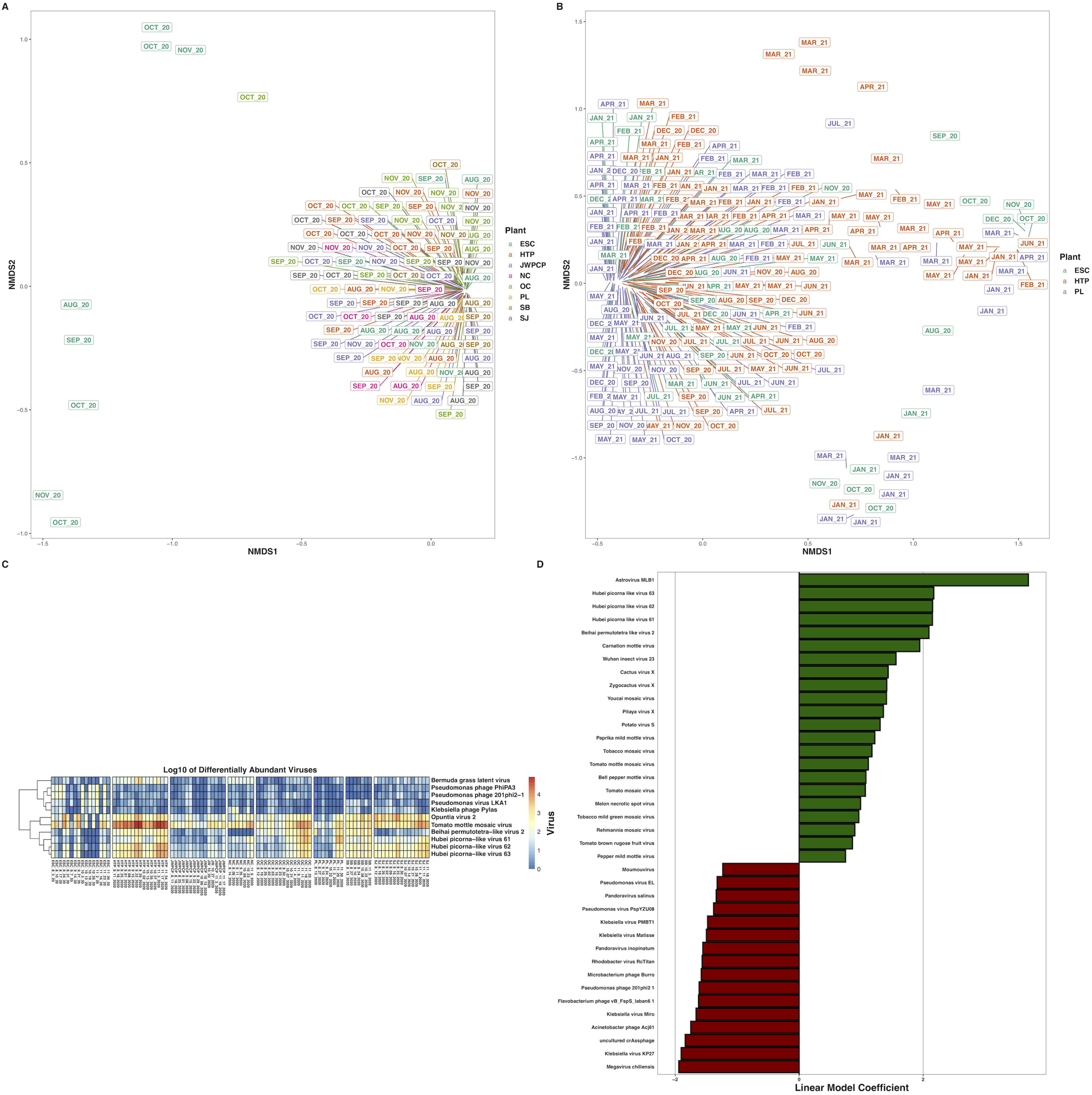
Nonmetric multidimensional scaling ordination of Bray-Curtis dissimilarities of viruses at greater than 0.01% relative abundance across A) all WTPs August – November 2020, and B) ESC, HTP, and PL across August 2020 – August 2021. C) Heatmap of the log_10_-transformed counts of differentially abundant viruses across all WTPs August – November 2020. D) Bar plots indicating the viruses across ESC, HTP, and PL that changed over time. X-axes denote the linear model coefficient of each virus’s relationship to time.

The full-year samples showed significantly different alpha diversity between WTP (H_(2)_ = 55.4, P < 0.001) but not over time (t = 0.11, P = 0.91). Long-term beta diversity differed between WTPs (P = 0.005, R^2^ = 0.03), month (P = 0.001, R^2^ = 0.11), with no interaction between WTP and month (P = 0.34, R^2^ = 0.09), with significant batch effects (P = 0.001, R2 = 0.11), and changed significantly over time (t = 4.3, P < 0.001). When considering the proportional abundance of individual virus species over the year, 22 viruses increased 16 decreased, and 102 did not change over time (Fig. 5, supplemental file SF3, doi.org/10.5281/zenodo.6829029).

### 3.5 Metabolic pathway transcription

Across samples that successfully processed through HUMAnN3 (N = 252), we detected transcripts that mapped to 474 Metacyc metabolic pathways (Dryad: https://doi.org/10.7280/D11Q30). Most commonly, we found transcriptional activity from pathways such as nucleotide biosynthesis, ubiquitination, amino acid biosynthesis, and central carbon metabolism, while we also detected rarer pathways involved in the degradation of xenobiotics including toluene, atrazine, nitrobenzoate, and octane.

Metabolic transcript alpha diversity was not significantly different across WTPs from August – November 2020 (H_(7)_ = 4.8, P = 0.68) and did not change over time (t = 1.2, P = 0.222). Likewise, metabolic transcript beta diversity during this period was not different between WTPs (P = 0.18, R^2^ = 0.09), but slightly differed between months (P = 0.003, R^2^ = 0.07) with an interaction between month and WTP (P = 0.03, R^2^ = 0.24) (Fig. 6), with significant batch effects (P < 0.001, R^2^ = 0.06), but did not change over time (t = 0.9, P = 0.38). There were no differentially-expressed metabolic pathways across WTPs during this time period.

**Figure 6.**
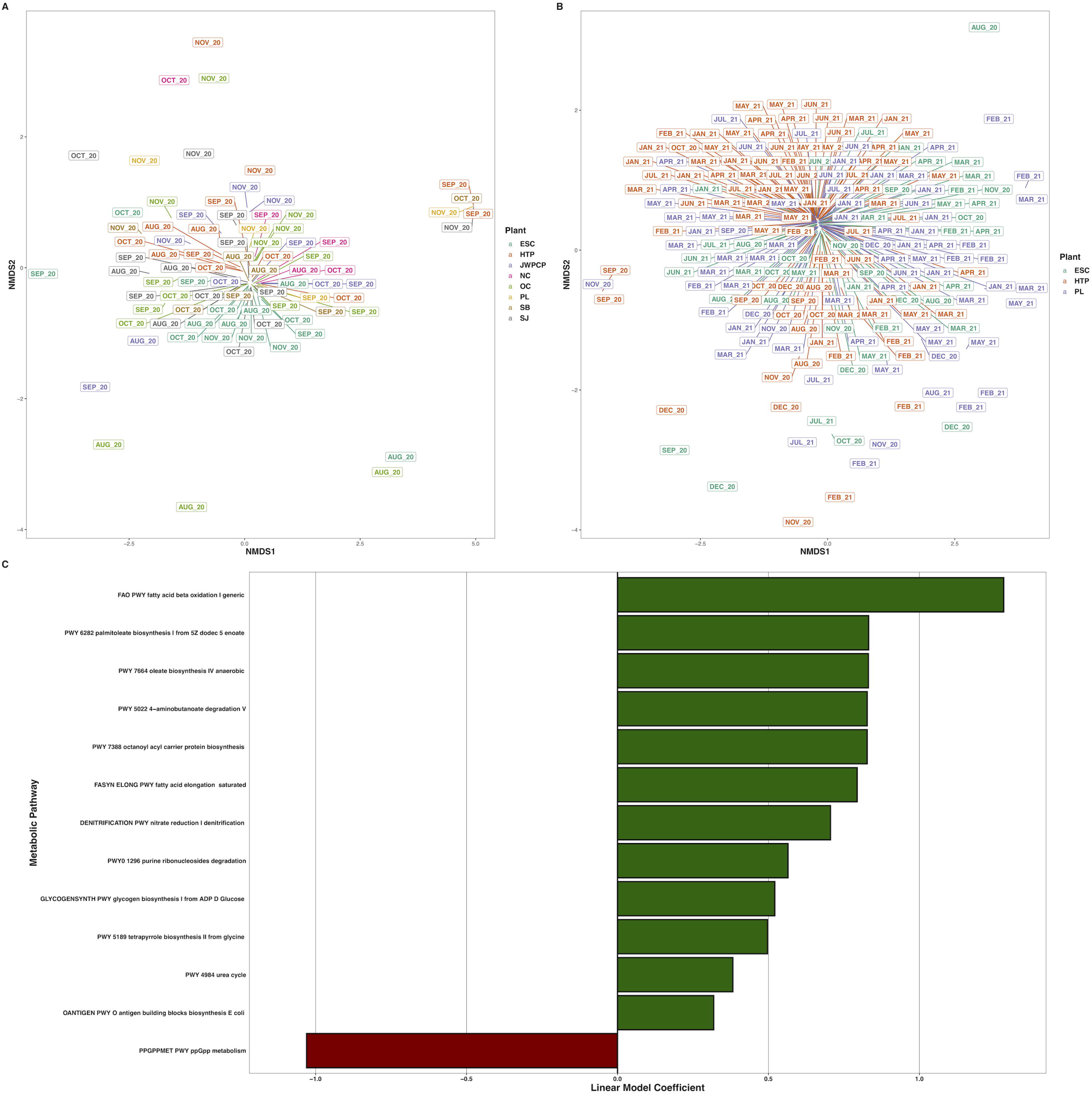
Nonmetric multidimensional scaling ordination of Bray-Curtis dissimilarities of metabolic pathway transcripts per million across A) all WTPs August – November 2020, and B) ESC, HTP, and PL across August 2020 – August 2021. C) Bar plots indicating the metabolic pathway at greater than 0.01% relative abundance across ESC, HTP, and PL that changed over time. X-axes denote the linear model coefficient of each metabolic pathway’s relationship to time.

Across the full year, transcript alpha diversity differed between WTPs (H_(2)_ = 14.4, P < 0.001), but not over time (t = 1.3, P = 0.19). Beta diversity slightly differed between WTPs (P = 0.008, R^2^ = 0.02), month (P < 0.001, R^2^ = 0.17), with an interaction between WTP and month (P = 0.005, R^2^ = 0.12) with significant sequencing batch effects (P = 0.002, R^2^ = 0.04), and did not change longitudinally (t = 0.8, P = 0.41). The transcription of few metabolic pathways had a significant association with time, as only 12 were positively, and one was negatively correlated, out of 205 pathways total (Fig. 6, supplemental file SF3, Zenodo: doi.org/10.5281/zenodo.6829029).

## 4. Discussion

Composite wastewater samples from Southern California over the year contained RNA transcripts derived from a wide diversity of microorganisms. To the best of our knowledge, our study representing a sewer shed of 16 million people is the most complete metatranscriptomic characterization of a large metropolitan region’s wastewater to date. Most notably, we show evidence of actively transcribed antimicrobial resistance (AMR) genes that encode resistance to a variety of commonly-administered antimicrobial drugs including macrolides, aminoglycosides, tetracycline and other AMR classes (Alcock et al., 2020). Likewise, we also show that bacterial transcription and RNA viral diversity differed between wastewater treatment plants (WTPs), and that sequencing wastewater RNA can be a useful tool for wastewater-based epidemiology (WBE) (Brumfield et al., 2022; Crits-Christoph et al., 2021; de Nies et al., 2021; Rothman et al., 2021; Xagoraraki and O’brien, 2020). Finally, we examined the total RNA pool and described metabolic pathway transcription to show that wastewater metabolism is largely consistent across WTPs and over time, but that there are slight signatures of geographical location (Gulino et al., 2020). Our results suggest that RNA sequencing is a viable tool to understand the complex matrix that wastewater represents and is useful in assaying the microbes associated with large populations.

### 4.1 Antimicrobial resistance transcription across Southern California wastewater

Wastewater is known to harbor an array of AMR genes, and several studies have sequenced and/or quantified many of these genes in wastewater (de Nies et al., 2021; Ju et al., 2019; Raza et al., 2022; Yin et al., 2021). Our study differs in that we demonstrate transcriptional activity through RNA-sequencing, rather than the genomic potential of the sampled organisms. We found a wide diversity of transcribed AMR genes in our data, including components of the multidrug efflux pumps adeFGH (Coyne et al., 2010) and its repressor acrS (Hirakawa et al., 2008), the gene tetQ (Nikolich et al., 1992), which encodes a ribosomal protection protein against tetracycline, *Staphylococcus aureus’s* multidrug efflux protein lmrS (Floyd et al., 2010), genes in the aminoglycoside resistance series aadA and aph(3”) (Ramirez and Tolmasky, 2010), and several variants of the glycopeptide resistance gene vanR (Courvalin, 2006). Many of these transcripts have been previously detected in WTPs, or in animals that resided in wastewater (Brumfield et al., 2022; Marcelino et al., 2019). Because we did not deplete rRNAs during library preparations, most of our bacterial transcripts were ribosomal RNAs. We detected rRNA mutations that confer macrolide resistance in the medically important taxa *Neisseria, Campylobacter, Salmonella, Helicobacter, Staphylococcus, Streptococcus, Klebsiella,* and many others. These genera (and subsequent AMR-resistant rRNAs) were ubiquitous in our samples and are often found in wastewater (Jankowski et al., 2022; Joseph et al., 2019; Ju et al., 2019). Our results indicate transcriptional evidence of widespread AMR activity, and we posit that this AMR presence is likely to be found in other wastewater catchments making metatranscriptomics useful for tracking AMR across wide areas (de Nies et al., 2021). The diversity of AMR genes in our samples differed between WTPs, and there were a few AMR genes differentially abundant between WTPs – mostly mutant rRNAs. This finding supports studies that show geographic differences between AMR (Raza et al., 2022; Yin et al., 2021), but there are likely other factors impacting the diversity of AMR, such as disease load in the served populations. Interestingly, we noticed a general increase over time in the proportional abundance of several transcripts from the major facilitator superfamily (MFS) and resistance-nodulation-cell division (RND) antibiotic efflux pumps - which are often implicated in multidrug resistance (Li and Nikaido, 2009) - along with beta-lactamases, and aminoglycoside/macrolide resistant rRNAs (Alcock et al., 2020). These data support studies showing an increase in antibiotic resistance (Ju et al., 2019) and the prevalence of AMR genes, but may also be impacted by seasonal changes in the waste stream (Yang et al., 2013). Likewise, as antibiotic use has risen during the COVID-19 pandemic (Langford et al., 2020; Rose et al., 2021), we may be observing a concurrent rise in AMR transcription in wastewater, although because our samples were solely collected during COVID-19, we are only able to speculate.

### 4.2 Viral ecology of Southern California wastewater

Plant-infecting tobamoviruses dominated the viromes of our samples regardless of source or time of year (Bačnik et al., 2020; Brumfield et al., 2022; Cantalupo et al., 2011; Crits-Christoph et al., 2021), although we also found substantial numbers of reads mapping to phages including crAssphage and assorted bacteriophages. While most known phages have DNA genomes, previous studies have identified phages in wastewater RNA (Crits-Christoph et al., 2021; Wilder et al., 2021). We may be detecting novel RNA viruses, or transcription of either DNA or RNA based phage genomes. Viral diversity differed when tested across all WTPs and over the full year, supporting studies that suggest geographical signatures of viruses in wastewater, and may be due to differences in human diet and viral excretion, along with disease dynamics in bacteria and/or eukaryotic hosts (Bibby and Peccia, 2013; Brumfield et al., 2022; Gulino et al., 2020). Likewise, several viruses were differentially abundant over time, which may be due to underlying infection trends or due to unknown seasonality effects (Brinkman et al., 2017; Kazama et al., 2016). While overall viral diversity was different between WTPs and changed over time, highly abundant viruses tended to be present in most samples, which may afford new targets in establishing microbial water quality or the detection of sewage pollution (Cao et al., 2015; Jiang et al., 2022; Kitajima et al., 2018). Similarly, we detected several human-infecting viruses (i.e. Norwalk Virus and SARS-CoV-2) which provides support for WBE efforts (Crits-Christoph et al., 2021; Nemudryi et al., 2020; Rothman et al., 2021, 2020; Xagoraraki and O’brien, 2020), and we suggest that RNA sequencing of wastewater should be used in conjunction with targeted and quantificational approaches to assist in passively monitoring diseases across large populations.

### 4.3 Bacterial ecology and metabolic pathways in Southern California wastewater

Similar to other studies, we detected transcripts from bacterial species in wastewater - mostly in the form of rRNA reads (de Nies et al., 2021; Joseph et al., 2019). Human pathogens were broadly represented in our data, including ESKAPE bacteria (*Enterococcus faecium, Staphylococcus aureus, Klebsiella pneumoniae, Acinetobacter baumannii, Pseudomonas aeruginosa,* and *Enterobacter* spp.), *Campylobacter jejuni, Salmonella* spp., *Helicobacter pylori, Haemophilus* spp., sexually transmitted infectious (STIs) agents, and bacteria commonly found in the environment. Much as with viruses, the bacterial profiles of WTPs were different, although many species were ubiquitous throughout the samples (Wu et al., 2019). There were also noticeable changes in the relative proportional transcript abundance over time, with many bacterial genera displaying a bimodal periodicity: Higher transcript abundance during Winter and Summer, and generally higher as time proceeded from August 2020 to August 2021. Other work has shown a distinct seasonality to the wastewater microbial community (Peces et al., 2022) - and our data supports this as well - although there are many other factors that can affect wastewater communities, such as pH, flux, dissolved oxygen, and detergents (Wu et al., 2019). Likewise, we recognize that our RNA extraction methods were harsh, and surely resulted in nucleic acid degradation, which likely affects the accuracy of our results (Schuierer et al., 2017). Non-ribosomal bacterial metabolism was apparent in our data with transcripts mapping to widely-conserved pathways such as nucleotide and amino acid biosynthesis and ubiquitination, with no pathways differing between WTPs or over time (Caspi et al., 2020). Collectively, our results suggest that sequencing bacterial species and their constituent metabolic pathways common to wastewater may be useful for monitoring disease through WBE, and that novel targets to assay microbial water quality may be possible.

## 5. Conclusion

In our opinion, this large-scale longitudinal dataset represents an unprecedented metatranscriptomic characterization of wastewater across a large population and region. We detected a wide diversity of transcribed AMR genes, suggesting that RNA sequencing is a powerful tool for WBE and may be useful in monitoring the spread and intensity of AMR. Within our study, we sequenced the viromes of a large portion of Southern California’s wastewater catchment area and show that plant-infecting viruses dominate the RNA viral fraction, which may have additional uses in detecting agricultural disease outbreaks. Similarly, we detected numerous human pathogens and observed changes in the relative proportions of these taxa, lending more credence to WBE as a vital component to public health and microbial water quality assays. We suggest that future transcriptomic studies target disease-causing taxa in wastewater to understand and refine WBE and its usefulness to human health more deeply.

## Acknowledgments

We thank the staff of the City of Escondido Hale Avenue Resource Recovery Facility, the City of Los Angeles Sanitation and Environment, Los Angeles County Sanitation District, Orange County Sanitation District, and City of San Diego Public Utilities for collecting influent.

This research was supported by the University of California Office of the President Research Grants Program Office (award numbers R01RG3732 and R00RG2814) awarded to JAR and KLW, and a Hewitt Foundation for Biomedical Research postdoctoral fellowship to JAR. This work was made possible, in part, through access to the Genomics High Throughput Facility Shared Resource of the Cancer Center Support Grant (P30CA-062203) at the University of California, Irvine, NIH shared instrumentation grants 1S10RR025496-01, 1S10OD010794-01, and 1S10OD021718-01, and access to computing resources from the UCI High Performance Cloud Computing Center.

## Supplemental file legends

Supplemental file SF1: Sample metadata.

Supplemental file SF2: ANCOM analyses outputs. Includes Wald scores, significance testing, and bacterial genus, virus, or ARO term being tested.

Supplemental file SF3: MaAsLin2 outputs. Includes linear model coefficients of proportional abundances over time, standard errors, sample N included and excluded, P-value, and Q-value for each bacterial genus, virus, ARO term, and HUMAnN3 pathway. Individual scatterplots for each term being tested are available on Zenodo at (doi.org/10.5281/zenodo.6829029).

